# Nardilysin regulates *Slc2a2* expression through ISLET1 recruitment to an evolutionarily conserved enhancer in pancreatic β-cells

**DOI:** 10.64898/2026.02.19.706474

**Authors:** Kiyoto Nishi, Narangerel Ganbaatar, Mikiko Ohno, Shinya Ikeda, Hirotaka Iwasaki, Mend Amar Batbaatar, Enkhjin Gansukh, Eiichiro Nishi

**Author notes:** Corresponding author: Eiichiro Nishi. Corresponding author: Kiyoto Nishi. Equally contributed (Kiyoto Nishi, Narangerel Ganbaatar).

## Abstract

GLUT2 (*Slc2a2*) is a key glucose transporter in pancreatic β-cells, and its reduced expression is closely linked to defective glucose-stimulated insulin secretion (GSIS) and diabetes. We previously reported that pancreatic β-cell–specific nardilysin (NRDC)-deficient mice (BetaKO) exhibit severe diabetic phenotype with defective GSIS and reduced *Slc2a2* expression in islets. However, because BetaKO mice also showed reduced MafA, a key upstream regulator of *Slc2a2*, along with an increased α-cell/β-cell ratio and other secondary changes that could influence GLUT2 levels, the mechanism by which NRDC regulates *Slc2a2* transcription remained unclear. Here, we demonstrate that NRDC controls *Slc2a2* expression in a β-cell autonomous and MafA-independent manner. By integrating publicly available ATAC-seq and ChIP-seq datasets, we identified four active enhancer regions around the murine *Slc2a2* locus, two of which are evolutionarily conserved in human islets. Luciferase assays revealed that NRDC selectively controls the activity of a conserved enhancer located 39k bp downstream of the *Slc2a2* transcriptional start site. Chromatin immunoprecipitation (ChIP) and re-ChIP assays further revealed that NRDC binds to this enhancer and is required for efficient recruitment of ISLET1, a transcription factor upstream of *Slc2a2*. These findings indicate that NRDC directly regulates *Slc2a2* in addition to MafA, highlighting multifaceted roles of NRDC in pancreatic β-cell gene regulation.

## 1. INTRODUCTION

Pancreatic β-cells secrete insulin in response to elevated blood glucose levels and play a central role in maintaining systemic glucose homeostasis. In rodent β-cells, glucose transporter 2 (GLUT2, *Slc2a2*) characterized by a high Km and high transport capacity is known to function as a key glucose sensor [1]. Under hyperglycemic conditions, GLUT2 rapidly imports glucose into β-cells, thereby promoting glucose metabolism and ATP production. The resulting increase in intracellular ATP closes ATP-sensitive potassium channels, induces membrane depolarization, and triggers insulin secretion, a process known as glucose-stimulated insulin secretion (GSIS). Consistent with this role, *Slc2a2*^−/−^ mice develop hyperglycemia and hypoinsulinemia [2], providing direct evidence that GLUT2 plays the essential role in glucose sensing and insulin secretion in β-cells. Furthermore, reduced *Slc2a2* expression and defective GSIS are frequently observed in diabetic animal models, underscoring the importance of GLUT2 in the pathophysiology of diabetes [3,4].

Although GLUT2 is the predominant glucose transporter in rodent β-cells, human β-cells rely mainly on GLUT1 and GLUT3. Nevertheless, GLUT2 is not dispensable in humans: homozygous mutations in GLUT2 cause neonatal diabetes [5], and single nucleotide polymorphisms in *SLC2A2* are associated with fasting glucose levels and susceptibility to type 2 diabetes [6–8]. These observations indicate that GLUT2 contributes to human β-cell function and diabetes risk despite its relatively lower abundance.

Given its critical role in β-cell physiology, GLUT2 expression is tightly regulated by multiple β-cell transcription factors, including PDX1 [9], NEUROD1 [10], MAFA [11], ISLET1 [12] and HNF1α [13]. Previous studies have identified several promoter/enhancer regions controlling mouse *Slc2a2* and human *SLC2A2* expression [14–16]. However, the precise mechanisms, particularly how these transcriptional factors are recruited to the regulatory elements, remain elusive.

Nardilysin (NRDC; N-arginine dibasic convertase) is a zinc-dependent peptidase of the M16 family that has emerged as a multifaceted regulator of energy metabolism. Global ablation of NRDC (*Nrd1*^−/−^ mice) leads to markedly impaired GSIS, accompanied by enhanced thermogenesis in brown adipose tissue (BAT) and increased systemic insulin sensitivity [17,18]. In subsequent studies, we demonstrated that NRDC in BAT regulates thermogenesis by controlling both the transcription [18] and the protein stability [19] of uncoupling protein 1 (UCP1). In white adipose tissue (WAT), NRDC regulates insulin sensitivity through pathways involving HIF1α and PPARγ [20] .

We previously demonstrated that pancreatic β-cell–specific deletion of NRDC (BetaKO) exhibit results in severely impaired GSIS and overt diabetes, at least in part due to reduced MafA expression [17]. During this analysis, we also observed marked downregulation of GLUT2, a well-established downstream target of MafA, in BetaKO islets. However, since these mice also display an increased number of α-cells as well as profound hyperglycemia, it remained unclear whether the reduction of *Slc2a2* reflected a direct regulatory role of NRDC or secondary consequences of altered islet composition and metabolic disturbance.

In the present study, we demonstrate that NRDC regulates *Slc2a2* expression in a MafA-independent manner. By integrating publicly available ATAC-seq and ChIP-seq datasets, we identified four candidate enhancer regions of *Slc2a2*, among which an evolutionarily conserved distal enhancer is selectively regulated by NRDC. Mechanistically, NRDC binds to the enhancer and facilitates the recruitment of ISLET1, thereby promoting *Slc2a2* transcription. These findings uncover a previously unrecognized enhancer-based mechanism by which NRDC controls β-cell glucose sensing and insulin secretion, providing new insights into the pleiotropic roles of NRDC in β-cell biology.

## 2. MATERIALS AND METHODS

### 2.1 Animal studies

*Nrd1*^−/−^ mice (Accession No. CDB0466K, https://large.riken.jp/distribution/mutant-list.html) were generated as previously described [21]. Pancreatic islets were isolated as previously described [17]. All animal experiments were conducted in compliance with relevant laws and institutional guidelines and were approved by the Animal Ethics Committee of Shiga University of Medical Science (approval numbers: 2021-2-5-H3; approval date: February 5, 2021). The study is reported in accordance with the ARRIVE guidelines (https://arriveguidelines.org).

### 2.2 Cell culture, lentivirus production and infection

MIN6 cells [22] were grown in Dulbecco’s Modified Eagle Medium supplemented with 10% fetal bovine serum and antibiotics. Lentivirus production and infection are performed as previously described [23]. Briefly, lentiviral vectors expressing mouse *Mafa* or BLOCK-iT™ miR RNAi (Life Technologies) targeting *Nrd1* is produced with In-Fusion HD cloning kit (Takara Bio, Japan; #639648) and a lenti vector pBOBI [24]. The vectors and lentivirus-packing plasmids (PMDL/REV/VSVG) were transfected into HEK293T cells to produce lentiviral particles. MIN6 cells were infected with lentiviral particles in the presence of 8mg/mL polybrene.

### 2.3 ATAC-seq and ChIP-seq data analysis

Peaks of mouse ATAC-seq (GSM3271265) and ChIP-seq (GSM3271247, GSM7447002, GSM6594224, GSM6594225) data were obtained through ChIP-atlas [25], where data are aligned to mouse reference genome with Bowtie2[26] and peaks are called with MACS2 [27]. The data were visualized with Integrative Genomics Viewer [28]. Human ATAC-seq and ChIP-seq data [15] are obtained and visualized by using Islet Regulome Browser (https://pasqualilab.upf.edu/app/isletregulome) [29]. Sequence conservation of human and mice was analyzed with the evolutionarily conserved region (ECR) browser (https://ecrbrowser.dcode.org) [30].

### 2.4 Luciferase assay

Luciferase assay was performed as previously described [17]. Briefly, PicaGene Promoter Vector 2 (PGV-P2; Toyo Ink, Japan) with or without the insertion of mouse *slc2a2* promoter or enhancers was transiently transfected into MIN6 cells. pRL-TK renilla luc (Promega) was co-transfected with PGV-P2 to normalize transfection efficacy. Luciferase activity was quantified by the Dual-Luciferase Reporter Assay Kit (Promega). Primers used for subcloning are provided in Table S1.

### 2.5 Chromatin immunoprecipitation (ChIP) and Re-ChIP assay

Chromatin was prepared from MIN6 cells as previously described [18]. ChIP and re-ChIP assays were performed using ChIP-IT® Express ChIP Kits and Re-ChIP-IT® Express Magnetic Chromatin Re-Immunoprecipitation Kits (Active Motif). Antibodies against NRDC (mouse monoclonal antibody #2E6) and Islet-1 (ab20670, abcam) were used. Primers used for qPCR are listed in Table S1.

### 2.6 Quantitative real-time PCR (qRT-PCR) analysis

qRT-PCR was performed as previously described [23]. The results were standardized for comparison by measuring the level of β-actin (*Actb*) mRNA in each sample. Primers used for qPCR are listed in Table S1.

### 2.7 Statistics

Results are expressed as mean ± SEM. Student’s t test, one-way ANOVA with post hoc Tukey-Kramer test or two-way ANOVA with post hoc Sidak test was performed using GraphPad Prism 10.

## 3. RESULTS

### 3.1. NRDC regulates *Slc2a2* in a β-cell–autonomous and MafA-independent manner

We previously reported that β-cell–specific NRDC-deficient mice (BetaKO) exhibit markedly reduced *Slc2a2* expression in pancreatic islets, accompanied by severe diabetes and impaired GSIS. Although these findings established that NRDC is required for maintaining *Slc2a2* expression *in vivo*, it remained unclear whether the reduction of *Slc2a2* was a consequence of systemic metabolic disruption or an intrinsic defect within β-cells. To directly address β-cell autonomy, we performed NRDC knockdown in mouse pancreatic β-cell line MIN6. As shown in Fig. 1A, NRDC knockdown significantly suppressed *Slc2a2* expression. Furthermore, pancreatic islets from systemic NRDC-deficient mice (*Nrd1*^−/−^), which exhibit normal fasting blood glucose levels and α-cell / β-cell ratios [17], showed significantly reduced *Slc2a2* expression levels (Fig. 1B). These data suggest that NRDC regulates *Slc2a2* expression in a cell-autonomous manner.

**Fig. 1.**
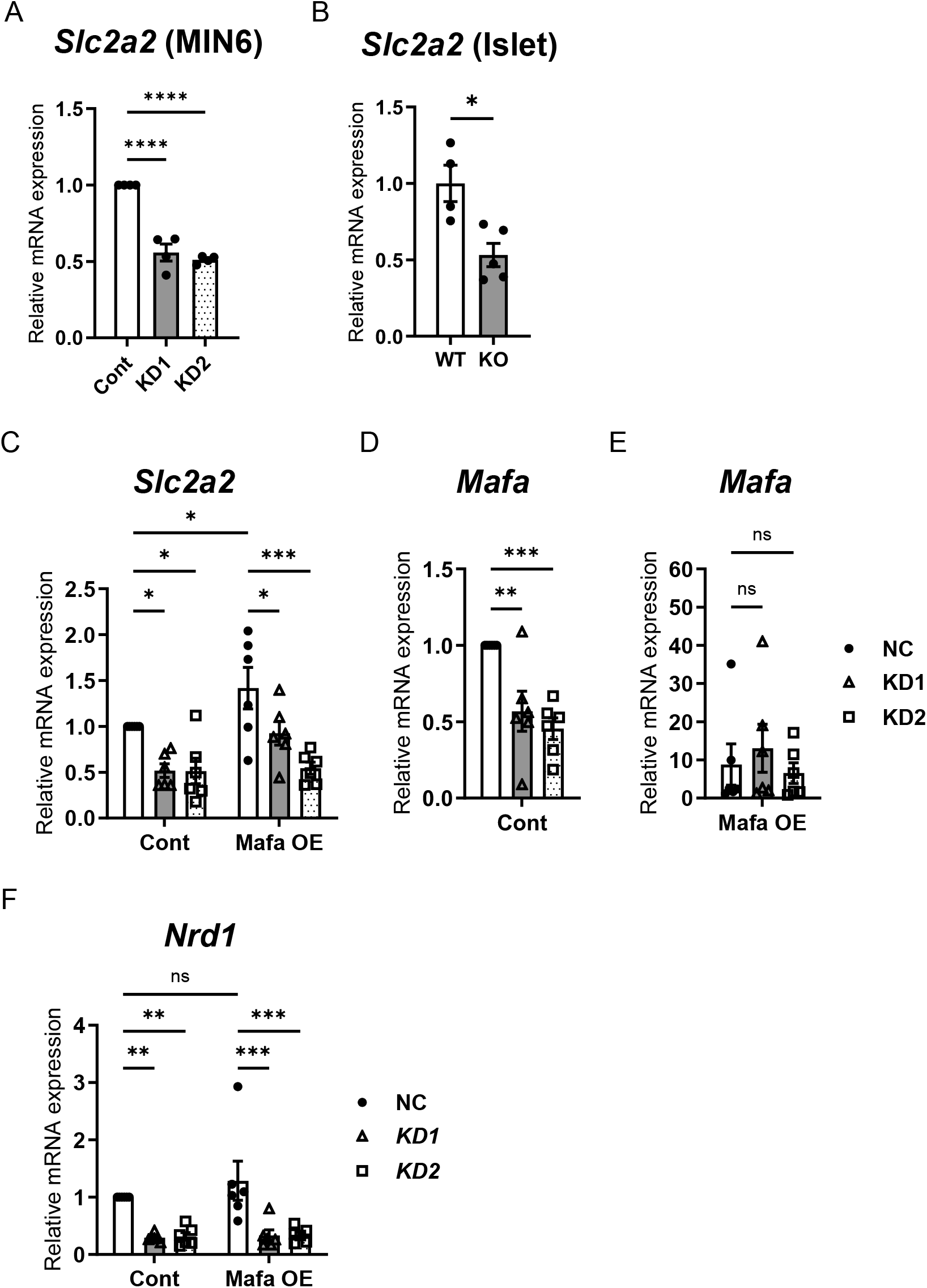
NRDC regulates *Slc2a2* in a β-cell–autonomous and MafA-independent manner. (A) Relative mRNA levels of *Slc2a2* in MIN6 cells with *Nrd1* knockdown (KD1, KD2) or control cells (n=4). (B) Relative mRNA levels of *Slc2a2* in pancreatic islets isolated from wild-type (WT) or *Nrd1*^-/-^ (KO) mice (n=4-5). (C-F): Relative mRNA levels of *Slc2a2* (C), *Mafa* (D-E), and *Nrd1* (F) in MIN6 cells with *Nrd1*-knockdown and *Mafa* Overexpression. Data expressed as mean ± SEM. ns, not significant, *: *p* < 0.05, **: *p* < 0.01, ***: *p* < 0.001, ****: *p* < 0.0001. *p* values were determined by unpaired Student’s t test (B), one-way ANOVA with post hoc Tukey-Kramer test (A, C-F), or two-way ANOVA with post hoc Sidak test (C, F).

As NRDC deficiency also lowers *MafA*, one of the key upstream activators of *Slc2a2* [17], we next examined whether *MafA* reduction accounts for the observed decrease in *Slc2a2*. Consistent with the regulatory role of MAFA over *Slc2a2, MafA* overexpression in MIN6 upregulated *Slc2a2* (Fig. 1C). Despite fully restoring *MafA* levels (Fig. 1D, E), *MafA* overexpression did not rescue the reduction of *Slc2a2* caused by NRDC knockdown (Fig. 1C). Importantly, *MafA* overexpression did not affect NRDC expression levels (Fig. 1F). Together, these findings demonstrate that NRDC regulates *Slc2a2* in a β-cell–autonomous and *MafA*-independent manner and that the transcriptional defect observed *in vivo* reflects an intrinsic requirement for NRDC in β-cells.

### 3.2. Identification of candidate enhancer regions regulating *Slc2a2* in MIN6 cells

To identify cis-regulatory elements through which NRDC may control *Slc2a2* expression, we surveyed publicly available ChIP-seq and ATAC-seq datasets from MIN6 cells using ChIP-Atlas [25]. Two prominent CCCTC-binding factor (CTCF) peaks were located 39k bp upstream and 49k bp downstream of the *Slc2a2* transcription start site (TSS) (Fig. 2A and Table S2) (GSM3271247) [16], delineating a 78kb area that likely constrains enhancer–promoter interactions. Within this region, analysis of ATAC-seq (GSM3271265) [16] and H3K27Ac ChIP-seq (GSM7447002, GSM6594224, GSM6594225) data [31,32] identified three potential candidate enhancer regions (ENH–16k, ENH+4k, and ENH+39k; Fig. 2A and Table S3,S4), fulfilling the following canonical criteria for active enhancers; (i) accessible chromatin regions detected by ATAC-seq and (ii) positive for H3K27ac. A previous study using islet rather than MIN6 datasets identified four potential enhancers (E5a, E5b, Ei and E3c) by similar criteria [14], and three of these (E5a, Ei, E3c) correspond directly to ENH–16k, ENH+4k, and ENH+39k, respectively. Because the MIN6 ATAC-seq dataset [7] showed a marginal but detectable accessibility peak at the remaining E5b region (located 2 kb upstream of the TSS), we also included this region as “ENH-2k” (Fig. S1A and Table S5).

**Fig. 2.**
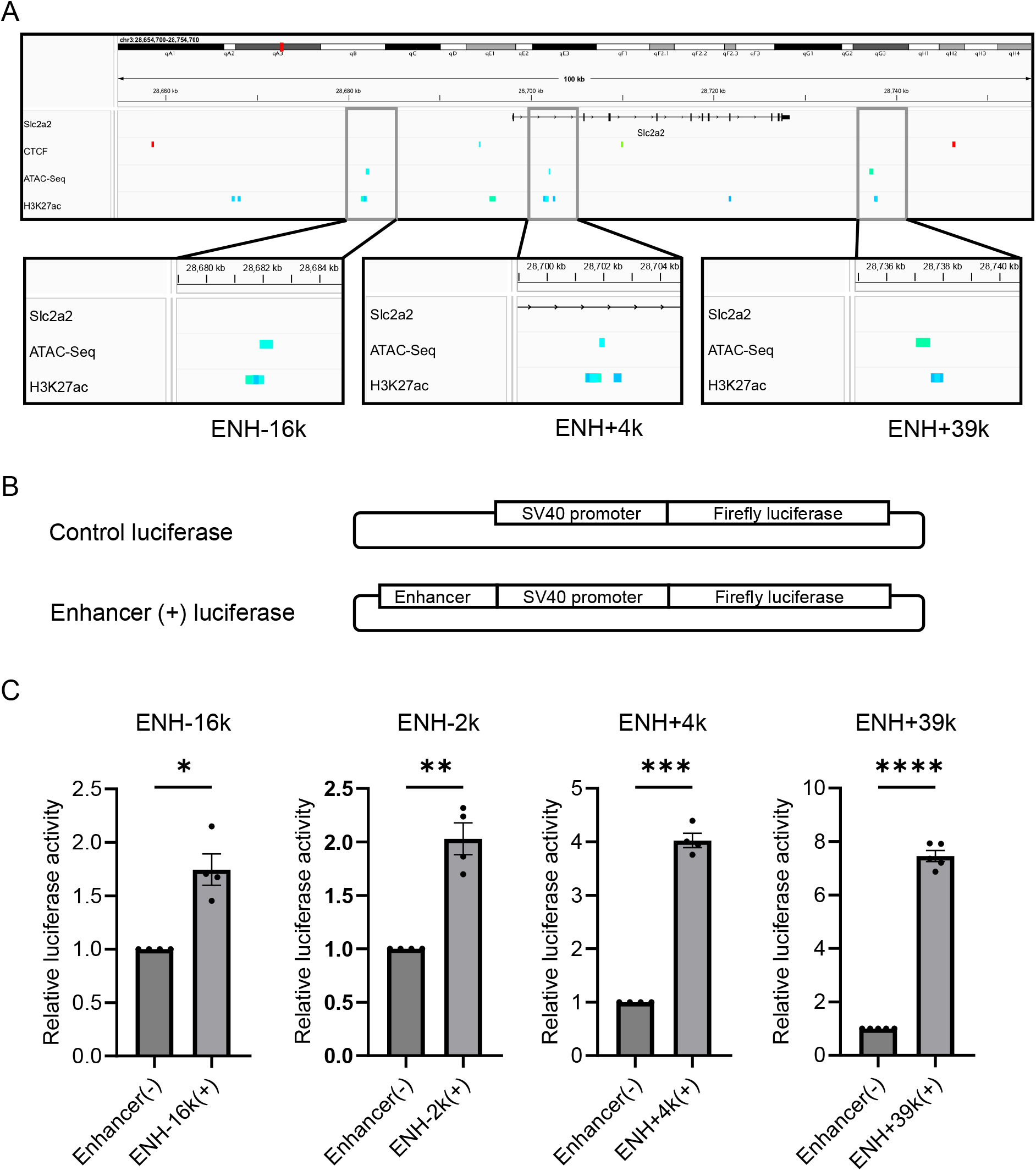
Identification of candidate enhancer regions regulating *Slc2a2* in MIN6 cells. (A) Peaks of ATAC-seq and ChIP-seq (CTCF and H3K27ac) around murine *Slc2a2* locus were visualized with Integrative Genomics Viewer. (B) Luciferase vectors used in Fig 2C. Possible enhancer sequences are inserted upstream of the SV40 promoter. (C) Luciferase assay using MIN6 cells transfected with reporter vectors with or without possible enhancer sequences (n=4-5). Data expressed as mean ± SEM. *: *p* < 0.05, **: *p* < 0.01, ***: *p* < 0.001. *p* values were determined by unpaired Student’s t test.

To assess whether these four candidate regions indeed exhibit enhancer activity, we cloned their ATAC-positive sequences into luciferase reporter vectors (ENH–16k, ENH-2k, ENH+4k, and ENH+39k) (Fig. 2B). For ENH+39k, we used a previously validated subfragment with demonstrated enhancer activity [12]. All four constructs containing enhancer regions significantly increased luciferase activity in MIN6 cells (Fig. 2C), confirming that each region functions as a bona fide enhancer for *Slc2a2*.

We next examined the evolutionary conservation of these enhancers. A comprehensive annotation of human islet enhancers [15] has identified eight enhancers within the 88-kb region flanked by two strong CTCF peaks surrounding the human *SLC2A2* promoter (Fig. S1B and Table S6), four of which are annotated as super-enhancers [33]. Using the Evolutionarily Conserved Region (ECR) browser (https://ecrbrowser.dcode.org) [30] we compared the murine and human loci and found that ENH-16k and ENH+39k show notable sequence conservation with the human enhancers (Fig. S1C).

Together, these analyses identified four murine *Slc2a2* enhancers, two of which are evolutionarily conserved in human islets and therefore strong candidates for mediating NRDC-dependent transcriptional regulation.

### NRDC regulates *Slc2a2* expression through an evolutionarily conserved enhancer

Given that ENH–16k, ENH–2k, ENH+4k, and ENH+39k all exhibited enhancer activity, we next asked which of these elements mediates NRDC-dependent regulation of *Slc2a2*. Luciferase assays using *Nrd1* knockdown and control MIN6 cells revealed a selective effect: only ENH+39k activity was significantly suppressed by *Nrd1* knockdown, whereas the activities of the other three enhancers were unaffected (Fig. 3A-D).

**Fig. 3.**
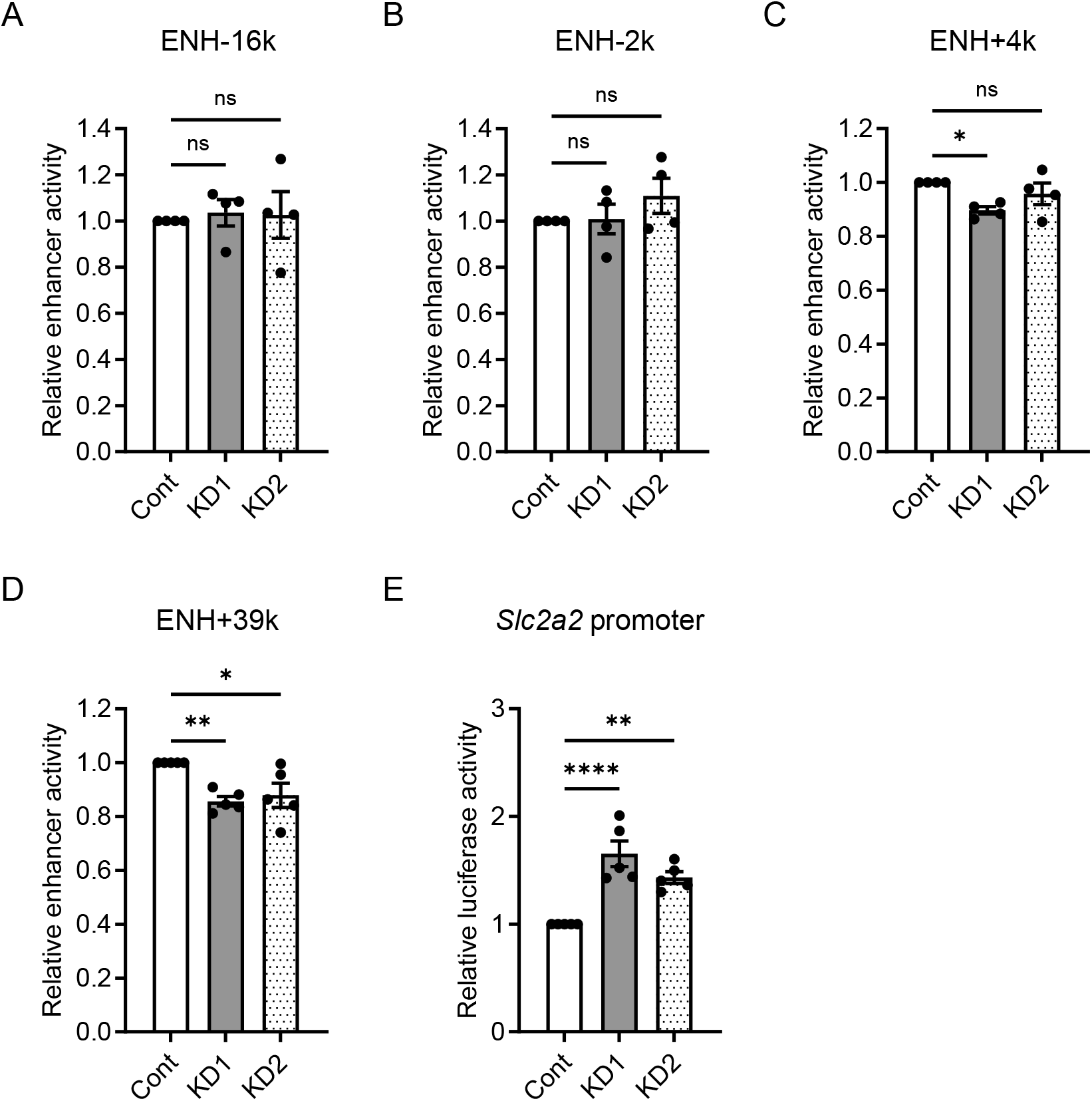
NRDC regulates *Slc2a2* expression through an evolutionarily conserved enhancer. (A-D) Luciferase assay using NRDC knockdown (KD1, KD2) or control (Cont) MIN6 cells transfected with reporter vectors with or without the enhancer sequences. Relative enhancer activity was defined as the luciferase activity of the enhancer (+) vector normalized to that of the enhancer (-) vector. (n=4-5). (E) *Slc2a2* promoter activity is assessed using NRDC knockdown (KD1, KD2) or control (Cont) MIN6 cells transfected with the reporter vector with *Slc2a2* promoter (n=5). Data expressed as mean ± SEM. ns, not significant, *: *p* < 0.05, **: *p* < 0.01, ***: *p* < 0.001. *p* values were determined by one-way ANOVA with post hoc Tukey-Kramer test.

Because *Slc2a2* transcription is also regulated through its promoter by multiple factors including MAFA [34], we examined whether NRDC influences promoter activity. Unexpectedly, *Nrd1* knockdown did not suppress Slc2a2 promoter activity; rather, promoter-driven luciferase activity was significantly upregulated (Fig. 3E). This finding indicates that the decrease in endogenous Slc2a2 expression upon NRDC deficiency cannot be explained by impaired promoter activation.

Together, these results indicate that NRDC regulates *Slc2a2* expression levels at least in part by modulating the activity of the evolutionarily conserved ENH+39k enhancer, rather than through the proximal promoter.

### NRDC facilitates the recruitment of ISLET1 to the *Slc2a2* ENH+39k enhancer

Previous studies have reported that several β-cell transcription factors, including PDX1, bind to ENH+39k enhancer and contribute to *Slc2a2* regulation. We first performed ChIP assays and confirmed the direct binding of NRDC to ENH+39k region (Fig. 4A). Among transcriptional factors known to interact with this enhancer, we focused on ISLET1 because it is the only transcription factor previously reported to physically interact with NRDC [17]. Consistent with a previous report [12], ISLET1 binding to ENH+39k was observed in MIN6 cells (Fig. S2A). These observations led us to hypothesize that NRDC cooperates with ISLET11 at the ENH+39k. ChIP-reChIP assays with anti-NRDC and anti-ISLET1 antibodies demonstrated that NRDC and ISLET1 co-occupy the ENH+39k region (Fig. 4B), supporting a model in which NRDC and ISLET1 act together at this regulatory element. To test whether NRDC influences the recruitment of ISLET1, we performed ChIP assays in *Nrd1* knockdown and control MIN6 cells. ISLET1 binding to ENH+39k was significantly reduced by NRDC knockdown (Fig. 4C).

**Fig. 4.**
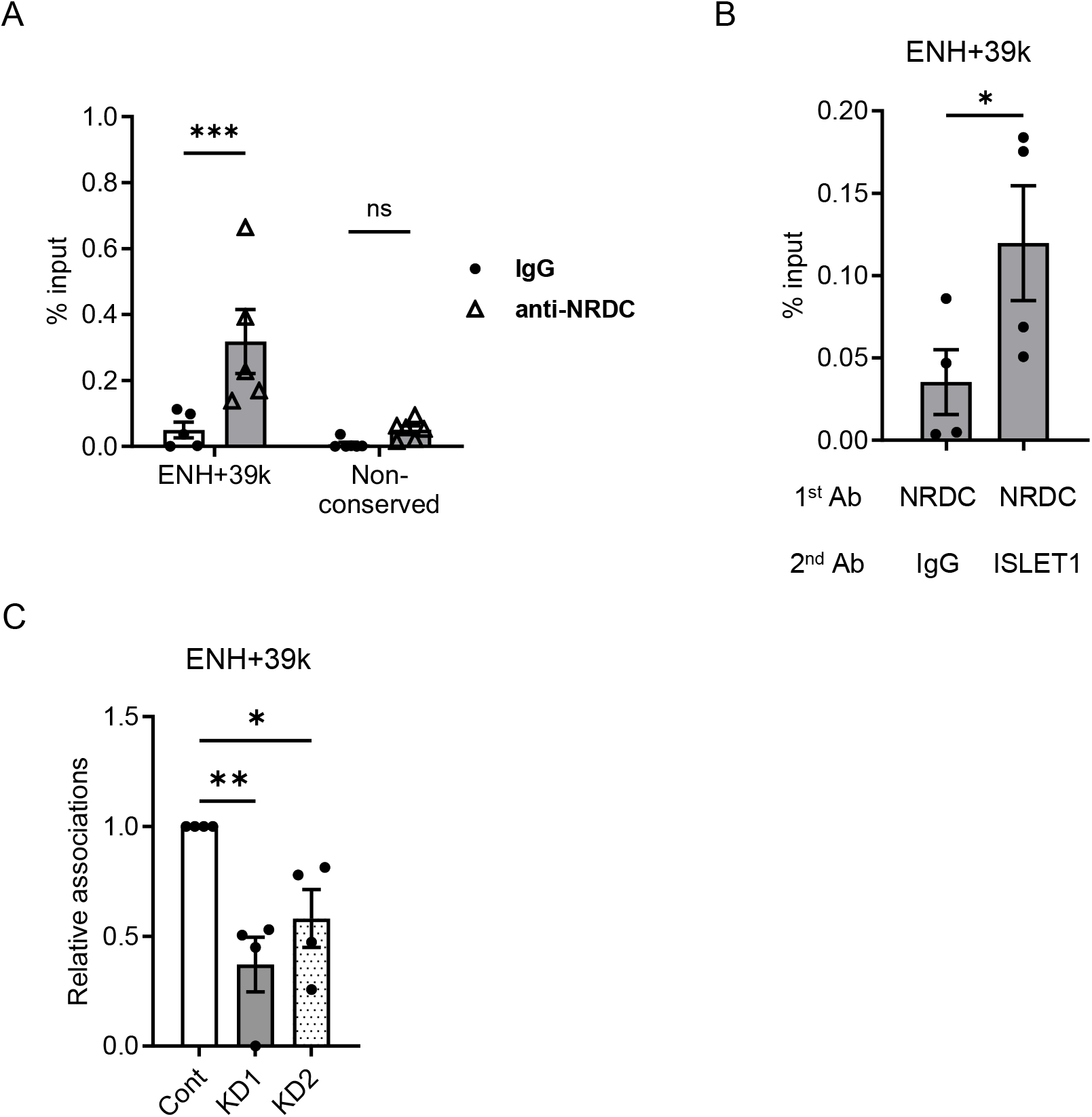
NRDC facilitates the recruitment of ISLET1 to the *Slc2a2* ENH+39k enhancer. (A) Chromatin immunoprecipitation (ChIP) assay with anti-NRDC antibody followed by quantitative RT-PCR targeting ENH+39k or Non-conserved region (-1678 to -1560 base pair relative to the TSS of Mafa). MIN6 cells are used and results were normalized to input DNA (n = 5-6). (B) ChIP/re-ChIP analysis of MIN6 cells with anti-NRDC antibody and anti-ISLET1 antibody followed by quantitative RT-PCR (qRT-PCR) targeting ENH+39k (n = 4). (C) ChIP assay with anti-ISLET1 antibody followed by quantitative RT-PCR using NRDC knockdown (KD1, KD2) or control (Cont) MIN6 cells. % input is normalized to that of control MIN6 cells (n=4). Data expressed as mean ± SEM. ns, not significant, *: *p* < 0.05, **: *p* < 0.01, ***: *p* < 0.001. *p* values were determined by unpaired Student’s t test (B), one-way ANOVA with post hoc Tukey-Kramer test (C), or two-way ANOVA with post hoc Sidak test (A).

Together, these findings demonstrate that NRDC promotes *Slc2a2* expression at least partly by facilitating ISLET1 recruitment to the conserved *Slc2a2* enhancer ENH+39k region, providing a mechanistic link between NRDC and GLUT2.

## 4. DISCUSSION

In this study, we elucidate a previously unrecognized regulatory mechanism by which NRDC maintains *Slc2a2* expression in pancreatic β-cells. While we previously demonstrated that NRDC regulates *MafA*, a key transcriptional activator of *Slc2a2*, the present study reveals that NRDC also controls *Slc2a2* expression through a *MafA*-independent pathway. By integrating publicly available ChIP-seq and ATAC-seq dataset with functional analyses in NRDC-knockdown MIN6 cells, we show that NRDC regulates *Slc2a2* expression through an evolutionarily conserved distal enhancer (ENH+39k). Mechanistically, NRDC directly binds this enhancer and facilitates the recruitment of ISLET1, establishing NRDC as a previously unappreciated co-regulator of enhancer-driven *Slc2a2* transcription in β-cells.

In addition to ISLET1[12], several transcription factors including MAFA, NEUROD1 and HNF1α have been shown to cooperatively activate *Slc2a2* expression in β-cells. As *Mafa* expression is suppressed by *Nrd1* knockdown, it remains possible that reduced ENH+39k activity in *Nrd1* knockdown cells is partially mediated through diminished *Mafa* levels. Thus, an important future direction will be to elucidate how the NRDC-ISLET1 axis intersects with the MAFA-NEUROD1-HNF1α network to fine-tune *Slc2a2* transcription.

Our identification of ENH–16k and ENH–2k as functional enhancers differs from previous reports in which these regions lacked enhancer activity[14]. This discrepancy may stem from methodological differences: whereas we analyzed narrow ATAC-seq–positive fragments, earlier studies employed larger genomic regions that may have included transcriptional repressive elements. Conversely, the enhancer activity of ENH+39k has been consistently reported across multiple studies [12,14,35], and this region is evolutionarily conserved as an enhancer (Fig. S1C). Together, these observations underscore ENH+39k as a core regulatory element governing *Slc2a2* expression in β-cells.

In addition to ENH+39k, the NRDC-regulated enhancer of *Mafa*[17] is also evolutionarily conserved in humans [36], raising the possibility that NRDC plays important roles in human β-cells. We have previously reported a family carrying a homozygous loss-of-function allele of NRDC, in which one affected individual with a homozygous truncating mutation was identified and clinically characterized. The presence of multiple long runs of homozygosity across the genome was consistent with distant parental relatedness. This individual exhibited a severe, progressive neurodevelopmental phenotype, including microcephaly, ataxia, and cerebral and cerebellar atrophy, highlighting an essential role for NRDC in neuronal maintenance [37]. Although no overt abnormalities in glucose homeostasis were reported, it is possible that defects in insulin secretion associated with NRDC deficiency remain clinically unapparent in the context of the overwhelming neurological phenotype.

Finally, our findings that NRDC also cooperates with ISLET1 not only in the regulation of *MafA* [17] but also of *Slc2a2*, raising the intriguing possibility that NRDC functions as a genome-wide cofactor for ISLET1-dependent gene regulation. Elucidating genome-wide co-occupancy of ISLET1 and NRDC by ChIP-seq or related approaches represents an attractive future direction. Given that ISLET1 plays critical roles in other organs including the heart, NRDC may similarly participate in ISLET1-dependent transcriptional programs in other organs, thereby contributing to diverse physiological and pathological processes.

## CRediT AUTHORSHIP CONTRIBUTION STATEMENT

Kiyoto Nishi: Writing – original draft, Visualization, Methodology, Investigation, Formal analysis, Data curation, Conceptualization, Funding acquisition. Narangerel Ganbaatar: Writing – original draft, Visualization, Methodology, Investigation, Formal analysis, Data curation. Mikiko Ohno: Writing – review & editing, Methodology, Investigation, Funding acquisition. Shinya Ikeda: Writing – review & editing, Methodology, Investigation. Hirotaka Iwasaki: Methodology, Investigation. Mend Amar Batbaatar: Methodology, Investigation. Enkhjin Gansukh: Methodology, Investigation. Eiichiro Nishi: Writing – review & editing, Supervision, Project administration, Methodology, Conceptualization, Funding acquisition.

## ACKNOWLEDGMENTS

MIN6 cells were generous gifts from Dr. Jun-ichi Miyazaki. We are grateful to M. Yoshida, R. Morinaga, M Sako and H. Iwai for their technical assistance, and T. Kita for continuous encouragement.

## DECLARATION OF GENERATIVE AI AND AI-ASSISTED TECHNOLOGIES IN THE WRITING PROCESS

During the preparation of this work the authors used ChatGPT and Google Gemini for English language editing, sentence shortening and minor corrections. After using these tools, the authors reviewed and edited the content as needed and take full responsibility for the content of the published article.

## DATA AVAILABILITY

Data will be made available on request.

