## Supplemental Figures and Tables for "Nardilysin regulates *Slc2a2* expression through ISLET1 recruitment to an evolutionarily conserved enhancer in pancreatic β-cells"

**SUPPLEMENTAL INFORMATION INVENTORY**

**Fig. S1-S2**

**Table S1-S6**

Fig. S1 related to Fig. 2

A

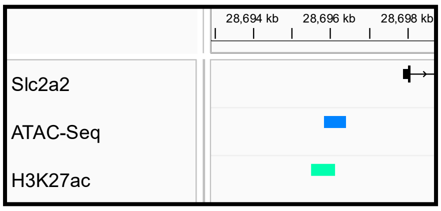


B

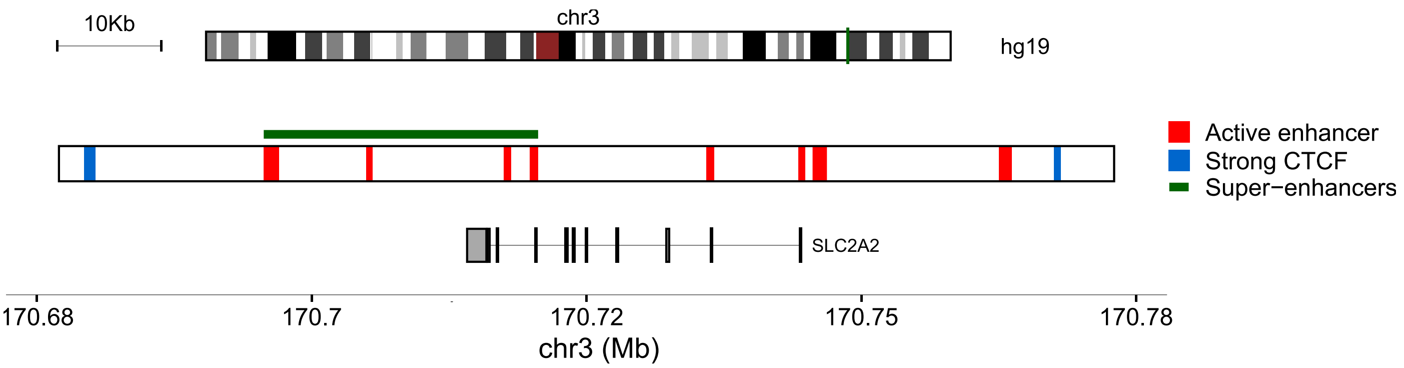


C

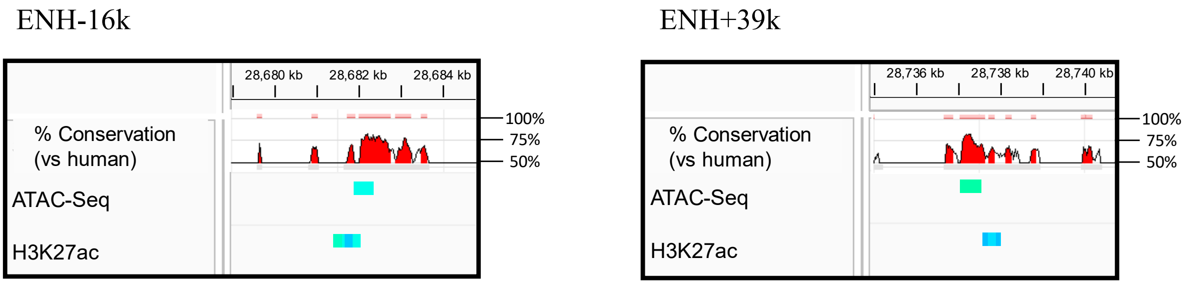


(A) Peaks of ATAC-seq and ChIP-seq (CTCF and H3K27ac) around murine *Slc2a2* promoter were visualized with Integrative Genomics Viewer.

(B) Strong CTCF peaks, active enhancers and super-enhancers around human *SLC2A2* locus were visualized with Islet Regulome Browser.

(C) ENH-16k and ENH+39k show notable sequence conservation with the corresponding human enhancers. Sequence conservation between human and mouse was quantified with the Evolutionarily Conserved Region (ECR) browser.

Figure S2 related to Figure 4

A

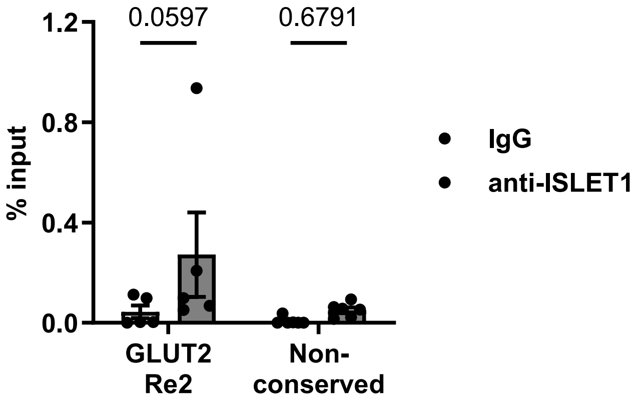


(A) Chromatin immunoprecipitation (ChIP) assay with anti-ISLET1 antibody followed by quantitative RT-PCR targeting ENH+39k or Non-conserved region (-1678 to -1560 base pair relative to the TSS of Mafa). MIN6 cells are used and results were normalized to input DNA (n = 5-6).

Table S1: List of Primers used for experiments.

| **Application** | **Forward** | **Reverse** |
|---|---|---|
| Subcloning mouse *Slc2a2* ENH-16k (chr3:28681781-28682365) into PGV-P2 | TCTATCGATAGGTACCACACTCCATAGTGCTTGACCA | GATCGCAGATCTCGAGATCTTTCTCTTATGAGTTGGTCTCTC |
| Subcloning mouse *Slc2a2* ENH-2k (chr3:28695825-28696398) into PGV-P2 | TCTATCGATAGGTACCCTCCACACTCACCCTGCA | GATCGCAGATCTCGAGACATTAATCTTTATAGCGTTCCCTTC |
| Subcloning mouse *Slc2a2* ENH+4k (chr3:28701841-28702174) into PGV-P2 | TCTATCGATAGGTACCTTTAGTCTGCAGGGACTC | GATCGCAGATCTCGAGTTCTAAGGGTCAGGATTTC |
| Subcloning mouse *Slc2a2* ENH+39k (chr3:28737100-28737399) into PGV-P2 | TCTATCGATAGGTACCGTCCAGTTCAAACTTGTTCCC | GATCGCAGATCTCGAGGTTACTTATTTGACAAAACAGAGGAG |
| Subcloning mouse *Slc2a2* promoter into PGV-P2 | CTAGCCCGGGCTCGAGCGACTCGATCCGTTGGCC | CCGGAATGCCAAGCTTTGTGTGGAATTGTCCTCTTAATCCA |
| qPCR mouse *Nrd1* | ATGGATGGCCTTTCCCTTG | CGCGAAGTTCAGCTTGTCAA |
| qPCR mouse *Slc2a2* | CATTCTTTGGTGGGTGGC | CCTGAGTGTGTTTGGAGCG |
| qPCR mouse *Mafa* | TTCAGCAAGGAGGAGGTCAT | CCGCCAACTTCTCGTATTTC |
| qPCR mouse *β-actin* | CTGACTGACTACCTCATGAAGATCCT | CTTAATGTCACGCACGATTTCC |
| ChIP qPCR mouse *Slc2a2* ENH+39k | CTTGTTCCCAAGTGACACCA | CTAACAGCAGGGAGCACACA |
| ChIP qPCR Non-conserved (-1678 to -1560 relative to the TSS of *Mafa*) | CCAGTTGCTTTTCACGGCCTC | CGGGGAGCCATTGGAATGTC |

Table S2: CTCF peaks from publicly available MIN6 ChIP-seq data set.

| Location | GEO Sample ID | SRA Experiment ID | score |
|---|---|---|---|
| chr3:28658435-28658715 | GSM3271247 | SRX4389955 | 1000 |
| chr3:28694251-28694463 | GSM3271247 | SRX4389955 | 252 |
| chr3:28709791-28710062 | GSM3271247 | SRX4389955 | 611 |
| chr3:28746117-28746495 | GSM3271247 | SRX4389955 | 1000 |

Table S3: Accessible chromatin regions detected by publicly available MIN6 ATAC-seq data set.

| Location | GEO Sample ID | SRA Experiment ID | score |
|---|---|---|---|
| chr3:28681846-28682350 | GSM3271265 | SRX4389961 | 270 |
| chr3:28701867-28702088 | GSM3271265 | SRX4389961 | 249 |
| chr3:28737012-28737534 | GSM3271265 | SRX4389961 | 336 |

Table S4: H3K27Ac peaks from publicly available MIN6 ChIP-seq data set.

| Location | GEO Sample ID | SRA Experiment ID | score |
|---|---|---|---|
| chr3:28667169-28667508 | GSM7447002 | SRX20599680 | 216 |
| chr3:28667868-28668217 | GSM7447002 | SRX20599680 | 203 |
| chr3:28681343-28681805 | GSM7447002 | SRX20599680 | 309 |
| chr3:28681569-28682043 | GSM6594224 | SRX17640936 | 272 |
| chr3:28681652-28681870 | GSM6594225 | SRX17640937 | 198 |
| chr3:28695460-28696136 | GSM6594224 | SRX17640936 | 329 |
| chr3:28701328-28701926 | GSM6594225 | SRX17640937 | 297 |
| chr3:28701359-28701746 | GSM7447002 | SRX20599680 | 191 |
| chr3:28701522-28701842 | GSM6594224 | SRX17640936 | 245 |
| chr3:28702365-28702658 | GSM6594225 | SRX17640937 | 193 |
| chr3:28721639-28721880 | GSM6594224 | SRX17640936 | 176 |
| chr3:28737534-28738003 | GSM6594225 | SRX17640937 | 202 |
| chr3:28737547-28737982 | GSM7447002 | SRX20599680 | 188 |
| chr3:28737696-28737919 | GSM6594224 | SRX17640936 | 221 |

Table S5: Accessible chromatin regions detected by publicly available MIN6 ATAC-seq data set.

| Location | GEO Sample ID | SRA Experiment ID | score |
|---|---|---|---|
| chr3:28667531-28667786 | GSM3271265 | SRX4389961 | 173 |
| chr3:28681781-28682365 | GSM3271265 | SRX4389961 | 270 |
| chr3:28695825-28696398 | GSM3271265 | SRX4389961 | 129 |
| chr3:28701841-28702174 | GSM3271265 | SRX4389961 | 249 |
| chr3:28736990-28737714 | GSM3271265 | SRX4389961 | 336 |
| chr3:28746058-28746263 | GSM3271265 | SRX4389961 | 119 |

Table S6: CTCF-binding sites and active enhancer regions surrounding *SLC2A2* locus in human islets

| Location | Class |
|---|---|
| chr3:170679322-170680328 | Strong CTCF |
| chr3:170695631-170697014 | Active enhancer |
| chr3:170704970-170705542 | Active enhancer |
| chr3:170717487-170718146 | Active enhancer |
| chr3:170719860-170720597 | Active enhancer |
| chr3:170735924-170736634 | Active enhancer |
| chr3:170744273-170744869 | Active enhancer |
| chr3:170745567-170746872 | Active enhancer |
| chr3:170762505-170763689 | Active enhancer |
| chr3:170767527-170768156 | Strong CTCF |
